# A single-cell atlas of human glioblastoma reveals a single axis of phenotype in tumor-propagating cells

**DOI:** 10.1101/377606

**Authors:** Sören Müller, Elizabeth Di Lullo, Aparna Bhaduri, Beatriz Alvarado, Garima Yagnik, Gary Kohanbash, Manish Aghi, Aaron Diaz

## Abstract

Tumor-propagating glioblastoma (GBM) stem-like cells (GSCs) of the proneural and mesenchymal molecular subtypes have been described. However, it is unknown if these two GSC populations are sufficient to generate the spectrum of cellular heterogeneity observed in GBM. The lineage relationships and niche interactions of GSCs have not been fully elucidated. We perform single-cell RNA-sequencing (scRNA-seq) and matched exome sequencing of human GBMs (12 patients; >37,000 cells) to identify recurrent hierarchies of GSCs and their progeny. We map sequenced cells to tumor-anatomical structures and identify microenvironment interactions using reference atlases and quantitative immunohistochemistry. We find that all GSCs can be described by a single axis of variation, ranging from proneural to mesenchymal. Increasing mesenchymal GSC (mGSC) content, but not proneural GSC (pGSC) content, correlates with significantly inferior survival. All clonal expressed mutations are found in the GSC populations, with a greater representation of mutations found in mGSCs. While pGSCs upregulate markers of cell-cycle progression, mGSCs are largely quiescent and overexpress cytokines mediating the chemotaxis of myeloid-derived suppressor cells. We find mGSCs enriched in hypoxic regions while pGSCs are enriched in the tumor’s invasive edge. We show that varying proportions of mGSCs, pGSCs, their progeny and stromal/immune cells are sufficient to explain the genetic and phenotypic heterogeneity observed in GBM. This study sheds light on a long-standing debate regarding the lineage relationships between GSCs and other glioma cell types.

## Introduction

Glioblastoma (GBM) is the most aggressive cancer of the adult brain. GBM genetics have been studied extensively and yet targeted therapeutics have produced limited results. GBM remains essentially incurable.

GBMs have been classified into subtypes that have prognostic value based on gene expression (*1*). We and others have shown that GBMs contain heterogeneous mixtures of cells from distinct transcriptomic subtypes (*2*, *3*). This intratumor heterogeneity is at least partially to blame for the failures of targeted therapies.

GBM-propagating stem-like cells (GSC) have been identified that express genes matching the mesenchymal (e.g. *CHI3L1*(*YKL-40*), *CD44*) and proneural (e.g. *OLIG2*, *DLL3*) transcriptomic subtypes (e.g. *4*). However, the lineage relationship between proneural GSCs (pGSC) and mesenchymal GSCs (mGSCs) is unknown. Surprisingly little is known about the cellular progeny of GSCs in vivo and their interactions with their microenvironment. It is unclear if proneural and/or mesenchymal GSCs are sufficient to generate the heterogeneity observed in GBM.

We performed single-cell RNA sequencing (scRNA-seq) and whole-exome DNA sequencing (exome-seq) of specimens from untreated human GBMs. We integrated this with meta-analysis of sequencing data from The Cancer Genome Atlas (TCGA) (https://cancergenome.nih.gov/) and anatomical data from The Glioblastoma Atlas Project (GAP) (http://glioblastoma.alleninstitute.org/). Using immunohistochemistry (IHC) and automated image analysis of human GBM microarrays we validated phenotypes at the protein level.

We show that all GBM cells can be described by a single axis of gene signature, which ranges from proneural to mesenchymal. At the extremes of this axis reside stem-like cells which express canonical markers of mGSCs and pGSCs. All clonal expressed mutations in our specimens are found in the GSC subpopulations, with mGSCs having a higher representation of mutations. Our analysis shows that mGSCs, pGSCs, their progeny and stromal/immune cells are sufficient to explain the heterogeneity observed in GBM.

**Fig. 1.**
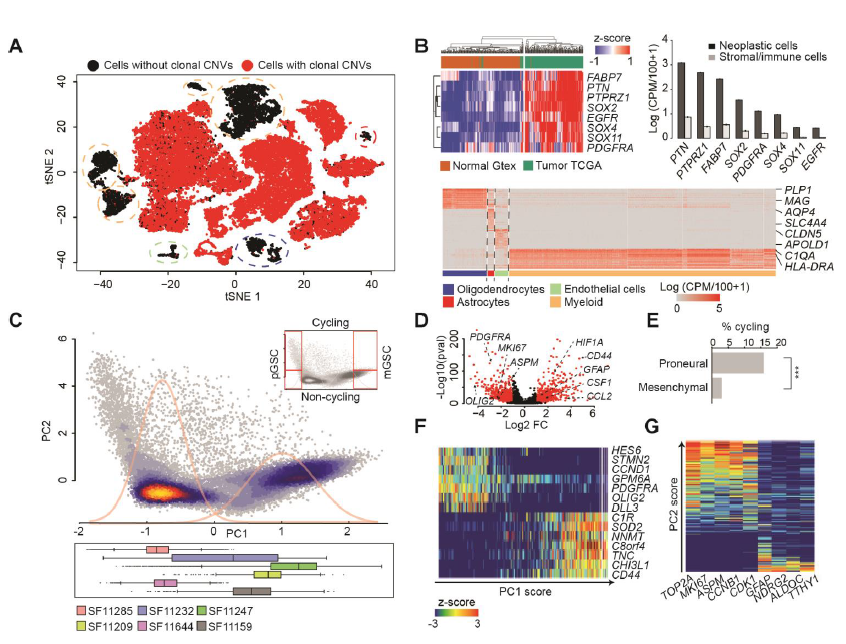
Single-cell sequencing reveals a single axis of variation in GBM cellular phenotypes. (A) A t-SNE plot of 32,151 10X cells from 6 patients. Cells are colored by the presence (red) or absence (black) of clonal CNVs. (B) (Top-left) Expression of GBM-enriched genes in IDH1-wildtype human GBMs from TCGA (n=144) and non-malignant human brain from Gtex (n=200). (Top-right) Average expression (+‐ SEM) of GBM marker-genes in cells classified as neoplastic (black) and non-neoplastic (grey). (Bottom) Heatmap of the 50 most specific genes (Wilcoxon rank-sum test) in clusters of non-neoplastic cells. (C) (Top) PCA of 25,899 10X neoplastic cells. Density curves of a Gaussian mixture model fit to PC1 sample scores are in gray. (Bottom) Distributions of cells from each patient along PC1. (D) Differentially expressed genes between PCA clusters (abs. log2 fold-change>1 and adj. p <0.001 in red). (E) Fractions of cycling cells, ***: Fisher p<0.001. (F-G) Expression of top-loading genes from PC1 (F) and PC2 (G) in single cells. Cells are sorted by PC1/2 sample score resp.

## Results

### Single-cell mRNA and bulk DNA profiling of human GBMs

We applied scRNA-seq to biopsies from 8 primary untreated human GBMs (table S1). Our goal was to profile both for cellular coverage (to survey cellular phenotypes) and for transcript coverage (to compare genetics). Therefore, we performed scRNA-seq on 6/8 samples via the 10X Genomics Chromium platform (10X) to obtain 3’ sequencing data for 37,196 cells. ScRNA-seq of the other 2/8 cases was done using the Fluidigm C1 platform (C1), which yielded full-transcript coverage for 192 cells. We incorporated 4 more published cases from our C1 pipeline, adding 384 cells (*3*). For 4 of the 10X cases and 4 of the C1 cases, the biopsies were minced, split and both scRNA-seq and exome sequencing (exome-seq) were performed (table S1). In total, scRNA-seq of 37,772 cells from 12 cases were used in this study.

We applied our pipeline for scRNA-seq quality control (*5*), quantification of expressed mutations (*3*, *6*), and cell-type identification (*7*, *8*). This identified 6,295 tumor-infiltrating stromal and immune cells based on expressed mutations, clustering and canonical marker genes (Fig. 1 A-B and Fig. S1A-D). We term the remaining 26,215 cells neoplastic, as they express clonal malignant mutations that have been validated by exome-seq (table S2). Only neoplastic cells were used for all subsequent analyses.

### The transcriptional phenotypes of GBM neoplastic cells can be explained by a single axis that varies from proneural to mesenchymal

An unbiased principal component analysis (PCA) revealed two patient-independent clusters of neoplastic cells (Fig. 1C, S1E-F; table S3). A differential-expression test between clusters identified canonical markers of the proneural (e.g. *PDGFRA*) and mesenchymal (e.g. *CD44*) subtypes as significant (table S4). Mesenchymal cells significantly over-express markers of response to hypoxia (e.g. *HIF1A*) and cytokines that promote myeloid-cell chemotaxis (e.g. *CSF1*, *CCL2*, *CXCL2*). However, mesenchymal cells do not express high levels of *MKI67* or other markers of cell-cycle progression. Conversely, proneural cells express high levels of *MKI67* as well as cyclin-independent kinase (Fig. 1D). We estimated the fraction of actively cycling cells using the Seurat package (*9*). By this metric, 15.6% of proneural cells are cycling compared to 2.9% of mesenchymal cells (Fig. 1E). Importantly, all our clinical specimens (assessed via either 10X or C1) contain cells of both phenotypes: proliferating proneural cells and mesenchymal cells with a quiescent, cytokine-secretory phenotype.

**Fig. 2.**
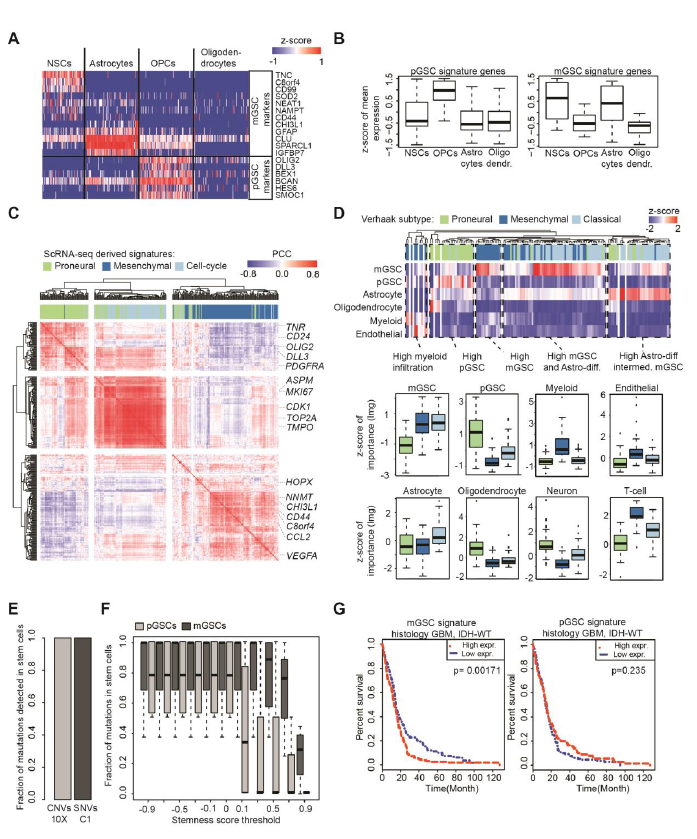
MGSCs and pGSCs explain the genetic and phenotypic heterogeneity of GBM. (A) Expression of mGSC and pGSC markers in single cells from non-malignant brain. (B) Z-scores of averages over pGSC and mGSC signature genes, compared across glial cell types. (C) Hierarchical clustering of Pearson correlations between mGSC, pGSC and cell-cycle genes in IDH1-wildtype GBM RNA-seq samples from TCGA (n=144). (D) Heatmap and boxplots of the relative contributions of predictor cell-types (rows) to the overall variance explained by a linear model fit to each TCGA sample. (E) Percentages of CNVs/SNVs expressed in GSCs in 10X/C1 data respectively, computed out of all mutations expressed in any patient’s cell that were validated by patient-matched exome-seq. (F) Percentages of SNVs (computed as in E) found in mGSCs and pGSCs for increasing stringency of stemness-score threshold. (G) Kaplan-Meier analysis comparing survival of IDH-wildtype GBMs from TCGA to average expression of the mGSC and pGSC gene signatures in patient-matched RNA sequencing.

Cells at the left and right extremes of principal component 1 (PC1) express high levels pGSC or mGSC markers (*4*) respectively, as indicated by PC1 gene loadings (Fig. 1F, S1D-E). We therefore interpreted the top-loading genes from each direction as representing pGSC and mGSC gene signatures. We used those signatures to score all cells for stemness, controlling for technical variation as previously described (*9*, *10*). Principal component 2 (PC2) correlates with cycling cells (Fig. 1G). Thus, while stemness correlates with cell cycle for both proneural and mesenchymal cells, more pGSCs than mGSCs express markers of cell-cycle progression, and at higher levels.

### GBM cells are stratified by differentiation gradients observed in gliogenesis

In our differential expression test we observed that proneural cells specifically express markers of the oligodendrocyte lineage (e.g. *OLIG2*, *SOX10*), while mesenchymal cells instead express markers of astrocytes (e.g. *GFAP*, *AQP4*). We evaluated the mGSC and pGSC gene signatures in scRNA-seq of glia from fetal and adult human brain (*11*, *12*). We found that pGSC-signature genes are enriched in oligodendrocyte progenitor cells. The mGSC signature, however, is comprised of genes expressed by neural stem cells as well as markers of astrocytes (Fig. 2A and 2B).

In addition to mGSCs and pGSCs, we find neoplastic cells (possessing clonal malignant mutations) that do not express stemness or cell-cycle signatures above background. Instead they express either markers of differentiated astrocytes (e.g. *ALDOC*) or differentiated oligodendrocytes (e.g. *MAG*, *MOG*). While the GSCs express high levels of positive WNT-pathway regulators, these more differentiated cells express high levels of WNT-pathway agonists (Fig. S2A). Thus, GBMs contain mesenchymal and proneural populations that align with astrocyte and oligodendrocyte differentiation gradients respectively.

### PGSCs, mGSCs, their differentiated progeny and stromal/immune cells explain the phenotypic heterogeneity observed in GBM

We found that our mGSC and pGSC gene signatures are co-expressed across TCGA datasets (Fig. 2C). While mGSC and pGSC signature genes are correlated among themselves, the mGSC and pGSC signatures are anti-correlated with each other. Moreover, the signature for cell-cycle progression obtained from PC2 more strongly correlates in TCGA data with the pGSC signature than the mGSC signature, consistent with our scRNA-seq data.

Using our scRNA-seq data and published scRNA-seq from human brain tissue (*11*, *12*) as a basis, we pooled reads across cells of the same type. This yielded data-driven profiles for mGSCs, pGSCs, astrocytes, oligodendrocytes, neurons, endothelial cells, myeloid cells and T-cells. We then used these cell-type signatures as predictors in a linear regression model. We fit our model to each TCGA GBM RNA-sequencing dataset individually (Fig. 2D). We found that samples of both the mesenchymal and classical Verhaak subtypes are enriched for mGSCs and depleted of pGSCs. While classical samples are distinguished by higher infiltration of astrocytes, mesenchymal samples contain high levels of infiltrating immune cells. Proneural samples are characterized by the highest levels of pGSCs, oligodendrocytes and neurons. In summary, the full spectrum of heterogeneity observed in TCGA GBM data can be explained by varying proportions of GSCs, their differentiated progeny and infiltrating stromal/immune content.

### GSCs explain their specimen’s genetic heterogeneity but only mGSC content is prognostic

We applied our pipeline for identifying single-nucleotide variants (SNVs) and megabase-scale copy-number variants (CNVs) to our exome-seq data (*3*, *6*). We restricted ourselves to mutations that occurred at a minimum of 10% variant allele frequency and identified cells in our scRNA-seq which expressed these mutations. For all patients, we found that all expressed, validated mutations are present in the GSCs (Fig. 2E and S3). We found that mGSCs possess a greater representation of mutations than pGSCs in all specimens. This result holds even if we increase the stringency in GSC assignment by thresholding the stemness score (Fig. 2F). Cox-regression analysis identifies the mGSC signature as correlating with significantly inferior survival in TCGA data. However, pGSC content is not prognostic (Fig. 2G).

### GSC niche localization and microenvironment interactions elucidated using reference atlases and quantitative IHC

We compared our mGSC, pGSC and cell-cycle signatures to RNA sequencing from the GAP (Fig. 3A-B). The GAP has annotated, micro-dissected and RNA sequenced GBM anatomical structures from human specimens. We found that mGSCs are enriched in hypoxic regions. PGSCs are enriched in the tumor’s leading edge and in regions of diffuse infiltration of tumor-adjacent white matter.

To infer GSC interactions with the microenvironment, we compared our scRNA-seq to a database of known receptor-agonist interactions. We annotated cognate pairs that were co-expressed by neoplastic and non-neoplastic cells from the same sample (table S5). We found that mGSCs compared to pGSCs express a greater diversity of surface receptors responsive to ligands expressed by stromal and immune cells (Fig. 3C).

To visualize and quantify the associations between GSCs, cell cycle and hypoxia, we performed IHC for CD44 (mGSCs), DLL3 (pGSCs), CA9 (hypoxia) and Ki67 (cell cycle) on GBM microarrays (Fig. 4A; table S6), as well as positive and negative control tissues (Fig. S4). We did not find any cycling CD44+ cells in our samples, while approximately 5% of DLL3+ cells expressed Ki67. On the other hand, CD44+ cells colocalized with CA9 at 2-fold greater frequency than DLL3+ cells (Fig. 4B). This dovetails with our findings in scRNA-seq, TCGA and GAP data, which show that pGSCs are more proliferative while mGSCs are enriched in hypoxic regions.

**Fig. 3.**
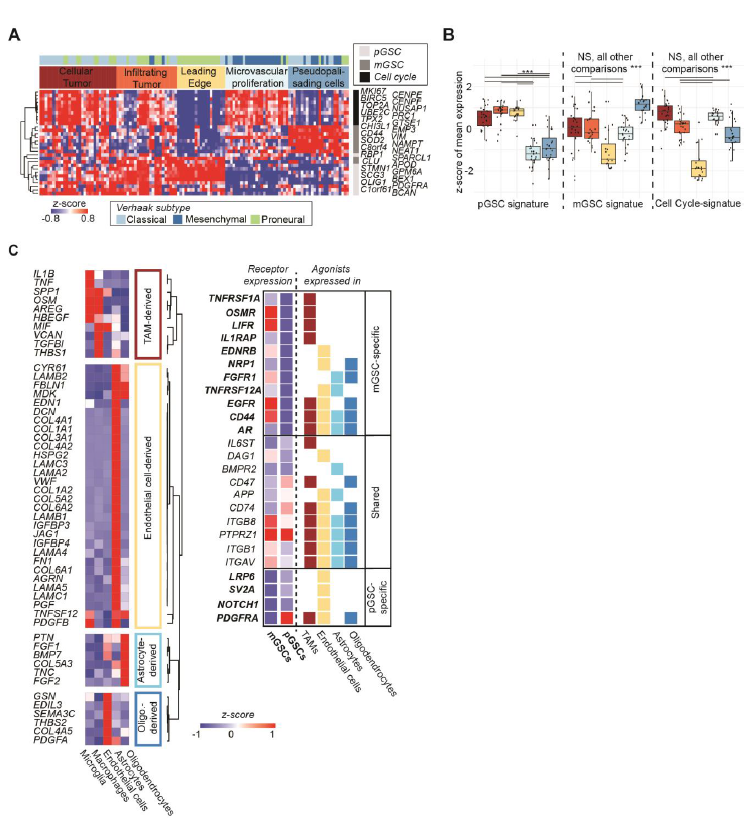
Niche interactions for mGSCs and pGSCs via sequencing. (A-B) MGSC, pGSC and cell-cycle signatures in GAP RNA-sequencing of GBM-anatomical structures. (C) Ligand expression in non-neoplastic cells (Left) and cognate receptor expression in neoplastic cells (Right).

## Discussion

It is known that an individual tumor may contain multiple GSC clones (e.g. *13*). We applied exome-seq and scRNA-seq to human tissues and found that all GBMs contain hierarchies of mesenchymal and proneural GSCs and their more differentiated progeny. However, the functional differences between GSC populations have not been fully determined. Murine GBMs can be separated into two cell populations that have different capacities for tumorigenicity and self-renewal (*14*). The Id1^high^/Olig2- and Id1^low^/Olig2+ populations found in the model of Barrett et al. match the expression signatures of mGSCs and pGSCs respectively. Moreover, the finding of Barrett et al. that Id1^high^ cells initiate tumors containing both Id1^high^ and Olig2+ cells, while tumors from Id1^low^/Olig2- cells do not generate Id1^high^ progeny, is consistent with our finding of a greater representation of clonal mutations in the mGSC population compared to pGSCs. Knowledge of GBM cell types, their lineage relationships and functional differences is needed to develop combination therapies that address intratumor heterogeneity.

**Fig. 4.**
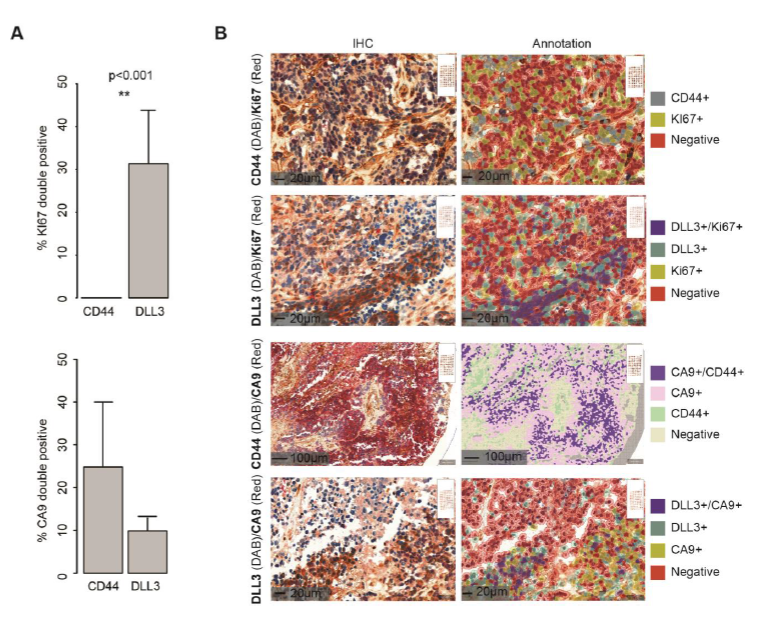
Niche interactions for mGSC and pGSC via IHC. (A) Percentages of CD44/DLL3 cells also positive for CA9/Ki67; **: Wilcoxon p<0.001. (B) Representative stains and image segmentation.

## Materials and Methods

### Tumor tissue acquisition and processing

We acquired fresh tumor tissue and peripheral blood from patients undergoing surgical resection for GBM. De-identified samples were provided by the Neurosurgery Tissue Bank at the University of California San Francisco (UCSF). Sample use was approved by the Institutional Review Board at UCSF. The experiments performed here conform to the principles set out in the WMA Declaration of Helsinki and the Department of Health and Human Services Belmont Report. All patients provided informed written consent. Tissues were minced in collection media (Leibovitz’s L-15 medium, 4 mg/mL glucose, 100 u/mL Penicillin, 100 ug/mL Streptomycin) with a scalpel. Samples dissociation was carried out in a mixture of papain (Worthington Biochem. Corp) and 2000 units/mL of DNase I freshly diluted in EBSS and incubated at 37 °C for 30 min. After centrifugation (5 min at 300 g), the suspension was resuspended in PBS. Subsequently, suspensions were triturated by pipetting up and down ten times and then passed through a 70-μm strainer cap (BD Falcon). Last, centrifugation was performed for 5 min at 300 g. After resuspension in PBS, pellets were passed through a 40-μm strainer cap (BD Falcon), followed by centrifugation for 5 min at 300 g. The dissociated, single cells were then resuspended in GNS (Neurocult NS-A (Stem Cell Tech.), 2 mM L-Glutamine, 100 U/mL Penicillin, 100 ug/mL Streptomycin, N2/B27 supplement (Invitrogen), sodium pyruvate).

### Fluidigm C1-based scRNA-seq

Fluidigm C1 Single-Cell Integrated Fluidic Circuit (IFC) and SMARTer Ultra Low RNA Kit were used for single-cell capture and complementary DNA (cDNA) generation. cDNA quantification was performed using Agilent High Sensitivity DNA Kits and diluted to 0.15–0.30 ng/μL. The Nextera XT DNA Library Prep Kit (Illumina) was used for dual indexing and amplification with the Fluidigm C1 protocol. Ninety-six scRNA-seq libraries were generated from each tumor/Cd11b + sample and subsequently pooled for 96-plex sequencing. cDNA was purification and size selection were carried out twice using 0.9X volume of Agencourt AMPure XP beads (Beckman Coulter). The resulting cDNA libraries were quantified using High Sensitivity DNA Kits (Agilent).

### 10X genomics-based scRNA-seq

Tissue was dissociated by incubation in papain with 10% DNAse for 30 min. A single-cell suspension was obtained by manual trituration using a glass pipette. The cells were filtered via an ovomucoid gradient to remove debris, pelleted, and resuspended in Neural Basal Media with serum at a concentration of 1700 cells/uL. In total, 10.2 uL of cells were loaded into each well of a 10X Chromium Single Cell capture chip and a total of two lanes were captured. Single-cell capture, reverse transcription, cell lysis, and library preparation were performed per manufacturer’s protocol. Sequencing for both platforms was performed on a HiSeq 2500 (Illumina, 100-bp paired-end protocol).

### Public data acquisition

Normalized counts from TCGA RNA-seq data were obtained from the Genomics Data Commons portal (https://gdc.cancer.gov/). Patients diagnosed as GBM and wild-type IDH1 expression (n = 44) were normalized to log2(CPM + 1) and used for analysis. TCGA GBM microarray and associated survival data were obtained from the Gliovis portal (*15*). Z-score normalized counts from regional RNA-seq of 122 samples from ten patients were obtained via the web interface of the Ivy GAP (http://glioblastoma.alleninstitute.org/) database.

### Exome-sequencing and genomic mutation identification

The NimbleGen SeqCap EZ Human Exome Kit v3.0 (Roche) was used for exome capture on a tumor sample and a blood control sample from each patient. Samples were sequenced with an Illumina-HiSeq 2500 machine (100-bp paired-end reads). Reads were mapped to the human grch37 genome with BWA (*16*) and only uniquely matched paired reads were used for analysis. PicardTools (http://broadinstitute.github.io/picard/) and the GATK toolkit (*17*) carried out quality score re-calibration, duplicate-removal, and realignment around indels. Large-scale (>100 Exons) somatic copy number variants (CNVs) were inferred with ADTex (*18*). To increase CNV size, proximal (< 1 Mbp) CNVs were merged. Somatic SNVs were inferred with MuTect (https://www.broadinstitute.org/cancer/cga/mutect) for each tumor/control pair and annotated with the Annovar software package (*19*).

### Single-cell RNA-sequencing, data processing and neoplastic-cell classification

Data processing of C1 data was performed as described previously (*3*). Briefly, reads were quality trimmed and TrimGalore! (http://www.bioinformatics.babraham.ac.uk/projects/trim_galore/) was used to clip Nextera adapters. HISAT2 (*20*) was used to perform alignments to the grch37 human genome. Gene expression was quantified using the ENSEMBL reference with featureCounts (*21*). Only correctly paired, uniquely mapped reads were kept. In each cell, expression values were scaled to counts per million (CPM)/100+1 and log transformed. Low-quality cells were filtered by thresholding number of genes detected at 1000 and at least 100,000 uniquely aligned reads. For 10x data we utilized CellRangeR for data pre-processing and gene expression quantification (version 1.3.6.).

### Classification of somatic mutations

The presence/absence of somatic CNVs in 10x data was assessed with CONICSmat (*6*). We retained CNVs with a CONICSmat likelihood-ratio test <0.001 and a difference in Bayesian Criterion >300. For each CNV we used a cutoff of posterior probability >0.5 in the CONICSmat mixture model to infer the presence/absence of that CNV in a given cell.

For the Fluidigm C1 data we utilized our previous CNV and SNV classifications (*3*). We retained only SNVs detected in exome-seq at >10% variant frequency in the tumor and <10% variant frequency in patient?matched normal blood. Cells expressing those tumor-restricted variant alleles were considered positive for the respective SNVs.

### Dimensionality reduction and calculation of stemness scores

The Seurat package was used for tSNE plots (*9*). PCA was done using R 3.4.2. The 1000 top genes with the highest biological variability were identified with the scran R package (*22*), for each patient. PCA was done using those genes that were amongst the 1000 most variable genes for at least three patients (757 genes). We defined the mGSC and pGSC gene sets as the top 15% of genes most strongly loading PC1, positively for mGSCs and negatively pGSCs. Stemness scores were calculated using these gene sets as input to the AddModuleScore function from the Seurat package.

### Deconvolution of TCGA RNA-sequencing data via linear models

To deconvolve GBM RNA-sequencing data from TCGA according to cell types learned from scRNA-seq, we first pooled scRNA-seq read-counts by cell type across mGSCs, pGSCs, non-malignant oligodendrocytes, astrocytes, neurons, TAMs, T-cells, and endothelial cells. The data used for this were our GBM scRNA-seq, as well as scRNA-seq of human fetal and adult non-malignant brain tissues (*11*, *12*). The resulting 8 count vectors were independently normalized to log2(CPM/10+1). We then fit a linear model to each TCGA RNA-sequencing dataset (also scaled to log2(CPM/10+1) using these vectors as predictors. We assessed the relative contribution of each predictor to the overall variance explained using the Lindeman, Merenda and Gold (lmg) method as implemented in the relaimpo R package (*23*).

### Immunohistochemistry

Immunohistochemistry (IHC) Optimization was performed on a Leica Bond automated immunostainer using conditions optimized for each antibody (table S5). Heat-induced antigen retrieval was performed using Leica Bond Epitope Retrieval Buffer 1 (Citrate Buffer, pH 6.0) and Leica Bond Epitope Retrieval Buffer 2 (EDTA solution, pH 9.0) for 20 minutes (ER2(20)). Non-specific antibody binding was blocked using 5% milk in PBST or Novocastra Protein Block (Novolink #RE7158). Positivity was detected using Novocastra Bond Refine Polymer Detection and visualized with 3’3 diaminobenzidine (DAB; brown) and alkaline phosphatase (AP; red). A Hematoxylin nuclear counterstain (blue) was applied. Duplex controls were performed to provide a reference of specificity of the selected primary antibody and secondary detection system. For the DLL3 and CA9 duplex, 2 sets of single-stained control tissues were used since the antigens of interest are not commonly co-expressed in normal tissue types. Image analysis of whole slide images (8 slides from GBM patients, 5 control slides) was performed using the Aperio software (Leica) and the ImageDx slide-management pipeline (Reveal Bio). All tissue and staining artifacts were digitally excluded from the reported quantification.

## Author Contributions and Notes

A.D. conceived of and designed the study. B.A., E.D., A.B. G.K and S.L. contributed to the scRNA-seq pipeline. M.A. provided biopsies via the UCSF Neurosurgery Tissue Core and contributed to the analysis of data. S.M. performed the bioinformatics analysis, under supervision of A.D. A.D. and S.M. wrote the manuscript, with input from all authors. All authors read and approved the final manuscript.

## Acknowledgments

We would like to thank Joanna Phillips and Anny Shai of the UCSF Neurosurgery Tissue Core who facilitated tissue acquisition.

## Ethics approval and consent to participate

Study protocols were approved by the UCSF Institutional Review Board (IRB# 11-06160). All clinical samples were analyzed in a de-identified fashion. All experiments were carried out in conformity to the principles set out in the WMA Declaration of Helsinki as well as the Department of Health and Human Services Belmont Report. Informed written consent was provided by all patients.

## Availability of data and material

The study data are available from the European Genome-phenome Archive repository, under EGAS00001002185, EGAS00001001900 and *(accession in process)*. Third-party data that were used in the study are available from the GlioVis portal (http://gliovis.bioinfo.cnio.es/), the gbmseq portal (http://gbmseq.org/), and The Glioblastoma Atlas Project (http://glioblastoma.alleninstitute.org/).

## Competing interests

None declared.

## Funding

This work has been supported by a Shurl and Kay Curci Foundation Research Grant, a UCSF Brain Tumor SPORE Career Development Award (P50-CA097257-13:7017), a Helen Diller Family Comprehensive Cancer Center/National Cancer Institute Cancer Center Support Grant (P30 CA 82103-18), and gifts from the Dabbiere Family and The Cancer League to A.D.

